# All eukaryotic SMC proteins induce a twist of -0.6 at each DNA-loop-extrusion step

**DOI:** 10.1101/2024.03.22.586328

**Authors:** Richard Janissen, Roman Barth, Iain F. Davidson, Michael Taschner, Stephan Gruber, Jan-Michael Peters, Cees Dekker

## Abstract

Eukaryotes carry three types of Structural Maintenance of Chromosomes (SMC) protein complexes, condensin, cohesin, and SMC5/6, which are ATP-dependent motor proteins that remodel the genome via DNA loop extrusion. SMCs modulate DNA supercoiling, but it has remained incompletely understood how this is achieved. Here we present a single-molecule magnetic tweezers assay that directly measures how much twist is induced by an individual SMC in each loop-extrusion step. We demonstrate that all three SMC complexes induce the same large negative twist (i.e., a linking number change Δ*L*k of -0.6 at each loop-extrusion step) into the extruded loop, independent of step size. Using ATP-hydrolysis mutants and non-hydrolysable ATP analogues, we find that ATP binding is the twist-inducing event during the ATPase cycle, which coincides with the force-generating loop-extrusion step. The fact that all three eukaryotic SMC proteins induce the same amount of twist indicates a common DNA-loop-extrusion mechanism among these SMC complexes.

## MAIN TEXT

SMC (Structural Maintenance of Chromosomes) protein complexes are essential in all organisms to ensure proper chromosome organization. In eukaryotes three SMC protein complexes exist: cohesin, condensin, and SMC5/6 (*1*). These SMC complexes share a common architecture (Fig. S1A) where two Smc subunits heterodimerize at their hinge and an intrinsically disordered kleisin subunit connects their ATPase domains, thus forming a ring structure. Cohesin and condensin associate with two HAWK (HEAT repeat-containing proteins Associated With Kleisins) subunits on the kleisin, while SMC5/6 carries two KITE (kleisin interacting winged-helix tandem element) on its kleisin and a third, Nse2, on the coiled coil of Smc5 (*2*). The three SMCs participate in a large range of diverse biological processes *in vivo* (*2*). From S-phase until metaphase, cohesin topologically embraces the two replicated sister chromatids (*3, 4*) while condensin compacts the chromatids *via* DNA loop extrusion in mitosis (*5, 6*) for subsequent chromosome segregation. During interphase, cohesin also folds genomic DNA into loops (*7–9*), which has been implicated in the regulation of transcription (*10*), recombination (*11, 12*), and the local separation of sister chromatids (*13*). SMC5/6 predominantly contributes to DNA repair (*14*), but its exact role remains elusive. Despite their varying roles, all three eukaryotic SMC complexes are able to extrude DNA into loops (*5, 8, 15–18*) *via* an ATP-dependent DNA loop extrusion mechanism that remains incompletely understood (*19*).

SMCs have been reported to change the amount of DNA supercoiling. Early biochemical reconstitutions of condensin from *Xenopus* egg extracts demonstrated that condensin generates positive DNA supercoiling (*20–22*), i.e., an additional twist deposited in DNA in the same right-handed direction as the double helix. Supercoiling is ubiquitously present in cells and contributes to the packaging of chromosomes and the regulation of genetic processes. Recent data have shown that condensing (*23*) and SMC5/6 (*24*) complexes are able to loop already supercoiled DNA and that cohesin also actively induce DNA supercoiling using ATP (*20, 23, 25*) and Davidson *et al*., 2024, bioRxiv). Condensin DCC has been associated with local negative supercoils at its high-occupancy binding sites of the X chromosome in *C. elegans* (*26*). Likely, DNA supercoiling and DNA loop extrusion stem from the same conformational changes during the ATPase cycle, suggesting that also SMC5/6 should exhibit a DNA supercoiling activity as well. Notably, DNA loop extrusion by condensing (*27*) and cohesion (*28*) occurs in large discrete steps of up to hundred base pairs each, which are generated by ATP binding to the SMC complex (*27*). It remains unknown how SMC proteins induce DNA twist, whether or how this mechanism is related to the loop-extrusion mechanism, and if the DNA twist directionality and degree of twist are comparable across the SMC family.

Here, we use a high-resolution magnetic tweezers assay to quantitatively examine, at the single-molecule level, the twist induced into DNA during loop extrusion by all three eukaryotic SMC complexes: human cohesin, yeast condensin, and yeast SMC5/6. We find that all three SMC complexes twist DNA negatively in each DNA loop-extrusion step, irrespective of the loop-extrusion step size. Notably, a large linking-number change (Δ*L*_k_ = -0.6) was resolved in each step of DNA loop extrusion by individual SMCs on single DNA molecules. Using ATP-hydrolysis mutants and non-hydrolysable ATP analogues, we show that ATP binding is not only the DNA extrusion step, but also the twist-generating event of the ATPase cycle. Strikingly, all three SMC complexes quantitatively induced the same amount of negative twist in the extruded DNA loop, indicating a conserved mechanism shared by all eukaryotic SMCs to generate supercoils that is an intrinsic property of the DNA loop-extrusion cycle.

### Assessing DNA twist that is induced at individual steps in DNA loop extrusion

With the objective to probe the degree of DNA twist induced by SMCs during single DNA loop extrusion steps, we turned to magnetic tweezers as this single-molecule technique has been proven to be able to resolve both single loop-extrusion steps of yeast condensin and human cohesion (*27, 28*), as well as minute changes in DNA twist (*29–32*). Torsionally constrained 3.6 kbp-long dsDNA molecules were tethered between a superparamagnetic 1 µm bead and an anti-digoxygenin-functionalized glass surface via handles on the end of DNA molecules containing either multiple biotin or digoxygenin labels, respectively (Methods). A pair of permanent magnets was mounted above the flow cell which allowed to exert a calibrated force on the surface-tethered DNA molecules (*33*) (Fig. 1A). Here, we applied a force of 0.3 pN, at which individual SMC-mediated loop extrusion steps can be measured (*27, 28*). Rotation of the magnet pair within the plane of the surface yielded a corresponding rotation of the magnetized bead. Since the attached DNA molecule is torsionally constrained, its linking number *L*_k_ changes proportionally with the rotation of the bead. The DNA extension is at its maximum when no rotation is applied, i.e., when Δ*L*_k_ = 0. Applying a low number of positive (negative) rotations overwinds (underwinds) and twists the DNA. Beyond a buckling point(*34*), twist is converted into writhe which absorbs the applied turns into plectonemic supercoils, resulting in the shortening of the DNA end-to-end extension (*29, 30, 32, 35*). Monitoring the DNA end-to-end extension versus positive and negative rotations thus generates a rotation curve which is symmetric around zero rotations (Δ*L*_k_ = 0) at low forces (≤ 0.6 pN, i.e., where DNA remains in its B-form (*29, 30, 32, 35*)) (Fig. 1A). Rotation curves were measured for all DNA molecules in the field of view as a reference before the introduction of SMCs into the flow chamber.

**Fig. 1.**
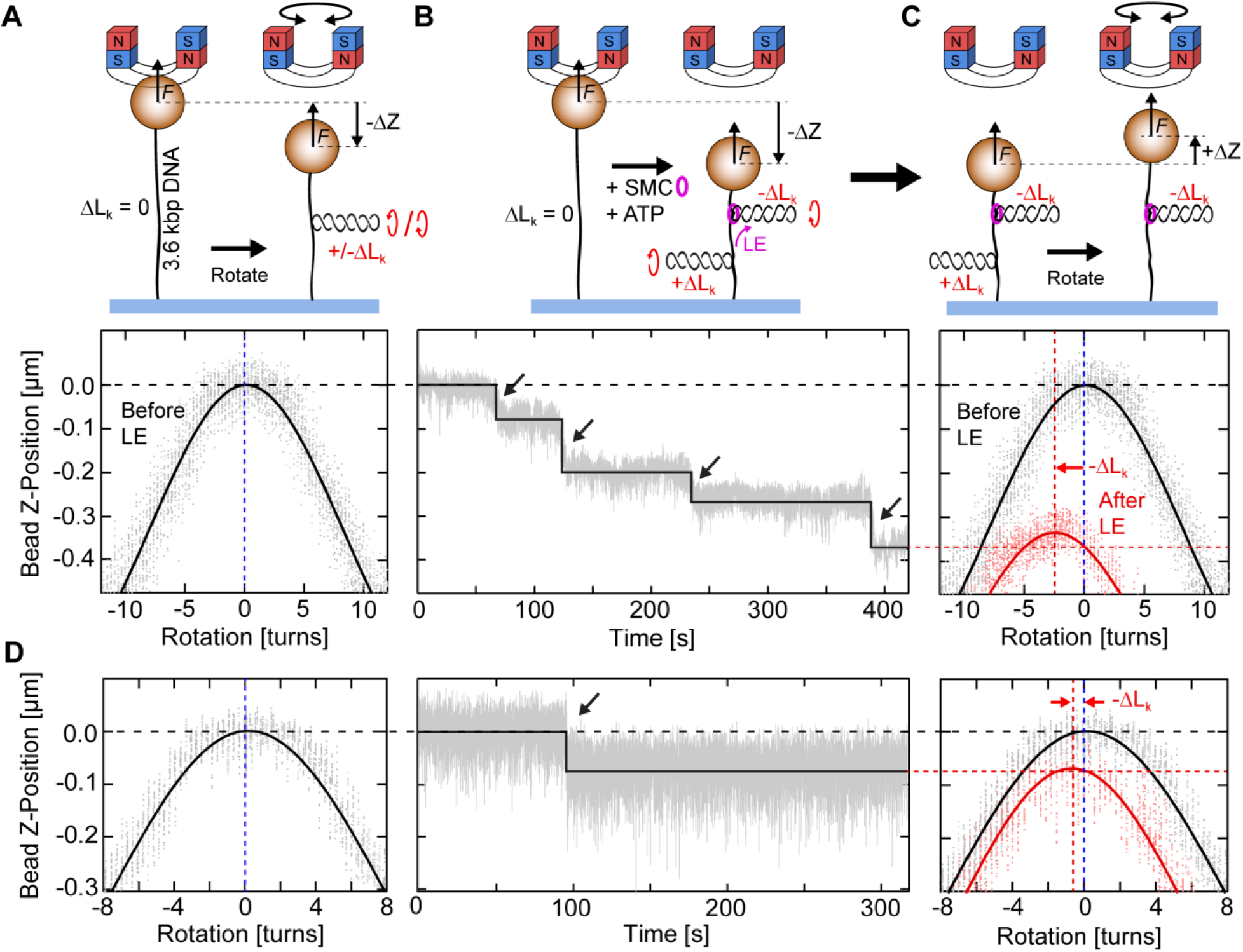
Measuring twist induced by single DNA-loop-extrusion steps by use of magnetic tweezers. (**A**) Assay schematic (top) and corresponding experimental data (bottom) of the DNA end-to-end extension as a function of magnet rotation for a torsionally constrained 3.6 kbp DNA molecule. At a constant force of 0.3 pN, the DNA extension is maximal when no external rotations are applied. Upon applying positive/negative rotations, the over/underwinding of the DNA changes the linking number *L*_k_ and leads to the formation of plectonemic supercoils, which, at this low force, are symmetric for both positive and negative coiling. Solid line is a Gaussian fit to the rotation curve data. (**B**) Assay schematic (top) and experimental data (bottom) which depict a representative trajectory of human cohesin (100 pM human cohesin, 250 pM NIPBL-Mau2) showing the step-wise DNA loop extrusion in the presence of 1 mM ATP at 0.3 pN. Black line shows fit result from a step-finding algorithm. During loop extrusion, human cohesin induces negative supercoils (-Δ*L*_k_) into the extruded DNA loop, while compensatory, positive supercoils (+Δ*L*_k_) form in the DNA molecule outside the loop. (**C**) Assay schematic (top) and experimental data (bottom) of DNA extension as a function of magnet rotation, similar to (A), conducted directly after the DNA loop extrusion experiment at 0.3 pN. The rotation curve after loop extrusion (red; line depicts Gaussian fit to the data) shows that the maximum DNA extension was shifted to negative magnet rotation compared to the initial rotation curve (black), which was caused by the uncoiling of the positive complementary supercoils (+Δ*L*_k_) that were formed in the DNA molecule outside the DNA loop. The degree of negative supercoils (-Δ*L*_k_) that are generated by human cohesin residing in the loop is thus directly measured from the shift of the rotation-curve maxima. Black arrows in the trajectories depict single steps. (**D**) Same as (A-C), for a trace showing only a single LE step (20 pM human cohesin with 50 pM NIPBL-Mau2 and 1 mM ATP). A lower amount of induced negative supercoils (-Δ*L*_k_) is observed, compared to (A-C).

This magnetic tweezers assay allows to resolve both the steps and the twist induced by individual SMCs during their loop extrusion activity. In a first series of experiments, human cohesin and ATP were flushed into the flow chamber while a high force of 7 pN was applied to the tethers to prevent compaction of the DNA molecules during flush-in by SMC-mediated DNA loop extrusion. When the flow was stopped and the force was quickly lowered to 0.3 pN, a stepwise decrease in the DNA end-to-end extension was observed (Fig. 1B). These can be attributed to discrete DNA loop extrusion steps as previously demonstrated for yeast condensin and human cohesion (*27, 28*). Directly after monitoring the steps for 10 minutes, another rotation curve was acquired at the same force (Fig. 1C). Interestingly, the peak of the rotation curve after loop extrusion (Fig. 1C, red curve) was systematically shifted to a lower bead height and to a negative number of rotations, compared to the reference rotation curve acquired before the extrusion of DNA by SMCs (Figs. 1A, 1C, black line). As the SMC reeled DNA into the loop, the DNA tether length decreased, explaining the decrease of the bead height (*27, 28*).

The shift of the rotation curve signals the twist that was inserted into the extruded loop by the SMC. To obtain the maximum DNA extension in the final rotation curve (Fig. 1C, red curve), negative rotations have to be externally applied to the DNA tether (outside the extruded loop) in order to remove the writhe. In other words, the DNA tether outside the loop was positively twisted after loop extrusion. Since the overall linking number of the entire torsionally constrained DNA molecule was conserved, the positive linking number change (Δ*L*_k_>0) that was measured in the non-looped DNA tether compensates the negative linking number change (Δ*L*_k_<0) that was generated by the SMC *inside* the extruded loop (Fig. 1C). In this way, our assay thus directly measures, at the single-molecule level, the induced twist that is generated into the extruded loop through one or more steps of the SMC. The magnitude of the induced twist Δ*L*_k_ is quantified as the difference of the peak positions of Gaussian fits to the rotation curves before and after loop extrusion (Methods), and it can be correlated to the number of downward steps that was observed in the time trace. A small but still negative shift between the rotation curves before and after the induction of loop extrusion could even be resolved for DNA molecules that showed only a single downward step (Fig. 1D). This, to our knowledge, constitutes the first direct demonstration that single SMC molecules negatively twist DNA, concomitant with individual loop extrusion steps.

### ATP-binding induces an equal amount of twist in each loop-extrusion step, irrespective of the step size

Using magnetic tweezers, we previously determined that ATP binding (i.e. not ATP hydrolysis) is the step-generating event during the DNA loop-extrusion cycle of yeast condensing (*27*). To test whether this is also the case for cohesin, as well as to assess whether the ATP-binding step is involved in DNA twisting, we used AMP-PnP, a non-hydrolysable ATP analogue, as well as an ATP-hydrolysis (EQ/EQ) deficient mutant during cohesin-mediated loop extrusion to obtain traces with a single loop-extrusion step. The DNA end-to-end extension exhibited only a single downward step (e.g. as in Fig. 1D) or repeated downward-reverse step combinations (Fig. S2A) for human cohesin in these conditions. No subsequent consecutive downward steps were observed, confirming that ATP binding is the step-generating event for human cohesin. Downward-reverse step combinations represent single loop extrusion steps which were followed by a reversal, possibly upon dissociation of DNA from one of the binding sites before the reeled in DNA was merged with the loop. Reversals were found to not contribute to the induced DNA twist (i.e. yielding Δ*L*_k_=0). The change in DNA linking number Δ*L*_k_ for traces with a single downward step (Fig. 2A), was -0.58 ± 0.16 (mean ± SD, N = 37).

**Fig. 2.**
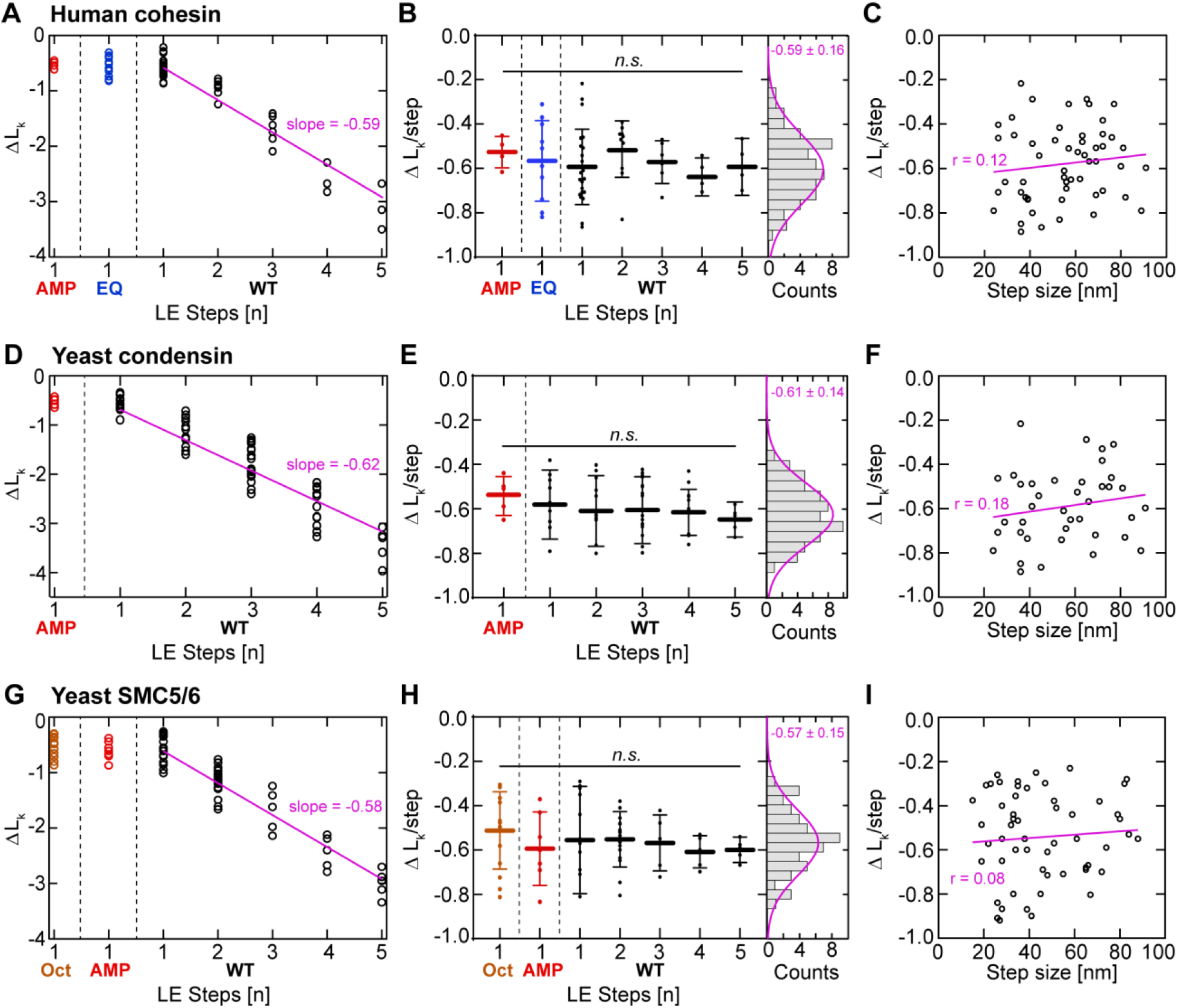
Induced DNA twist is constant for each loop extrusion step and step size-independent for all eukaryotic SMC complexes. (**A**) Linking number change Δ*L*_k_ versus the number of steps for WT human cohesin in the presence of AMP-PnP (red, N = 5), for EQ/EQ mutant of human cohesin in the presence of ATP (blue, N= 10), and for WT human cohesin in the presence of ATP (black, N = 63). Pink line is a linear fit without offset. (**B**) Same data as in panel (A), but displaying the linking number change per step (i.e. Δ*L*_k_ divided by the number of LE steps). Histogram on the right represents all data points (AMP-PnP, EQ, and WT) and is fitted by a Gaussian with -0.59 ± 0.16 Δ*L*_k_/step (mean ± SD). Statistical significance was assessed using a one-way analysis of variance (ANOVA) with a significance level *α* = 0.05 (95% confidence interval; n.s. = p > 0.05). (**C**) Linking number change versus the measured step size, for traces with only a single step (pooled from AMP-PnP, EQ, and WT experiments, N = 54). Linear fit and Pearson’s correlation coefficient are shown in pink. (**D-E**) As (A-C) but for yeast condensin (N = 5 for AMP-PnP, N=64 for WT, N = 39 in the right panel). (**G-I**) As (A) but for the yeast SMC5/6 hexamer (N = 15 for the SMC5/6 octamer [SMC5/6 with Nse5-Nse6], N = 7 for AMP-PnP, N = 56 for WT, N = 55 in the right panel).

By contrast, traces of wild type human cohesin in the presence of ATP often exhibited several consecutive downward steps (Fig. S2A). The change in linking number after cohesin-mediated loop extrusion was found to scale linearly with the number of loop extrusion steps (Fig. 2A). Division of the final linking number change by the number of loop extrusion steps in the trace showed that each step induced on average a DNA twist of Δ*L*_k_ = -0.59 ± 0.16 (mean ± S.D., N = 78; Fig. 2B). Strikingly, the linking number change was independent of the size of the loop extrusion step (Fig. 2C), as a similar -0.6 values were found for small (∼20 nm) steps and for large (∼100 nm) steps. This suggests that DNA twist is generated by local conformational changes within the SMC complex, independently of the amount of DNA that is fed into the loop.

### All eukaryotic SMCs induce Δ*L*k = -0.6 per step into the extruded DNA loop

All three eukaryotic SMC complexes – cohesin, condensin, and SMC5/6 – share the ability to extrude DNA into loops and they do so with very similar characteristics, viz., at a comparable extrusion rate and up to a comparable sub-pN stalling force (*5, 8, 15, 16, 28*). We therefore examined if their ability to twist DNA is also shared. To answer this question, we repeated magnetic tweezers experiments described above for human cohesin with budding yeast condensin (Figs. 2D-2F, S1B, S2B, Methods) and budding yeast SMC5/6 (Figs. 2G-2I, S1C, S2C, Methods, ref. (*36*)). We then analysed whether individual loop extrusion steps can be observed using torsionally unconstrained DNA and measured the size of these steps (Methods). At an applied force of 0.2 pN, we found that the median step size of SMC5/6 is ∼30 nm, while cohesin and condensin take steps of median size ∼40 nm (Fig. S1D). The step size decreased as the DNA tension is increased, a phenomenon observed for all eukaryotic SMC complexes (Fig. S1D and refs. (*27, 28*)).

Magnetic tweezers experiments on yeast condensin and SMC5/6 yielded very similar results as for human cohesin. In the presence of AMP-PnP, wild type yeast condensin and SMC5/6 showed only a single downward step and downward-reverse step combinations (Figs. S2B, S2C), while in the presence of ATP, multiple consecutive downward steps were observed (Figs. S2B, S2C). The SMC5/6 hexamer can be converted to an octameric version by the addition of the co-factor Nse5/6 (*37, 38*). Octameric SMC5/6 does not loop DNA (*16, 36*) but promotes salt-stable loading of SMC5/6 onto DNA, which is indicative of topological loading (*16, 37*). Interestingly, loop-extrusion traces of octameric SMC5/6 exhibited a single downward step or downward-reverse step combinations (but never multiple consecutive downward steps) (Fig. S2C), similar to traces of wild type SMCs in the presence of a non-hydrolyzable AMP-PnP or an ATP-hydrolysis deficient mutant (Fig. S2C and ref. (*27*), respectively). This suggests that Nse5/6 blocks ATP hydrolysis but not ATP binding to SMC5/6, consistent with an earlier report that ATP binding but not hydrolysis is required to topologically load SMC5/6 onto DNA (*37*).

As for cohesin, the linking number change induced by condensin and SMC5/6 hexamer was negative and scaled with the amount of loop extrusion steps, while the linking number change per step remained constant at a value of Δ*L*_k_ = -0.61 ± 0.14 and -0.57 ± 0.15, respectively (mean ± SD; Figs. 2E, 2H). These values were also independent of the size of the loop extrusion steps (Figs. 2F, 2I). Strikingly, all three SMC complexes thus induced a comparable amount of DNA twist per step of Δ*L*_k_ = -0.59 ± 0.02 (mean ± SD across the three SMCs, Fig. 3A).

**Fig. 3.**
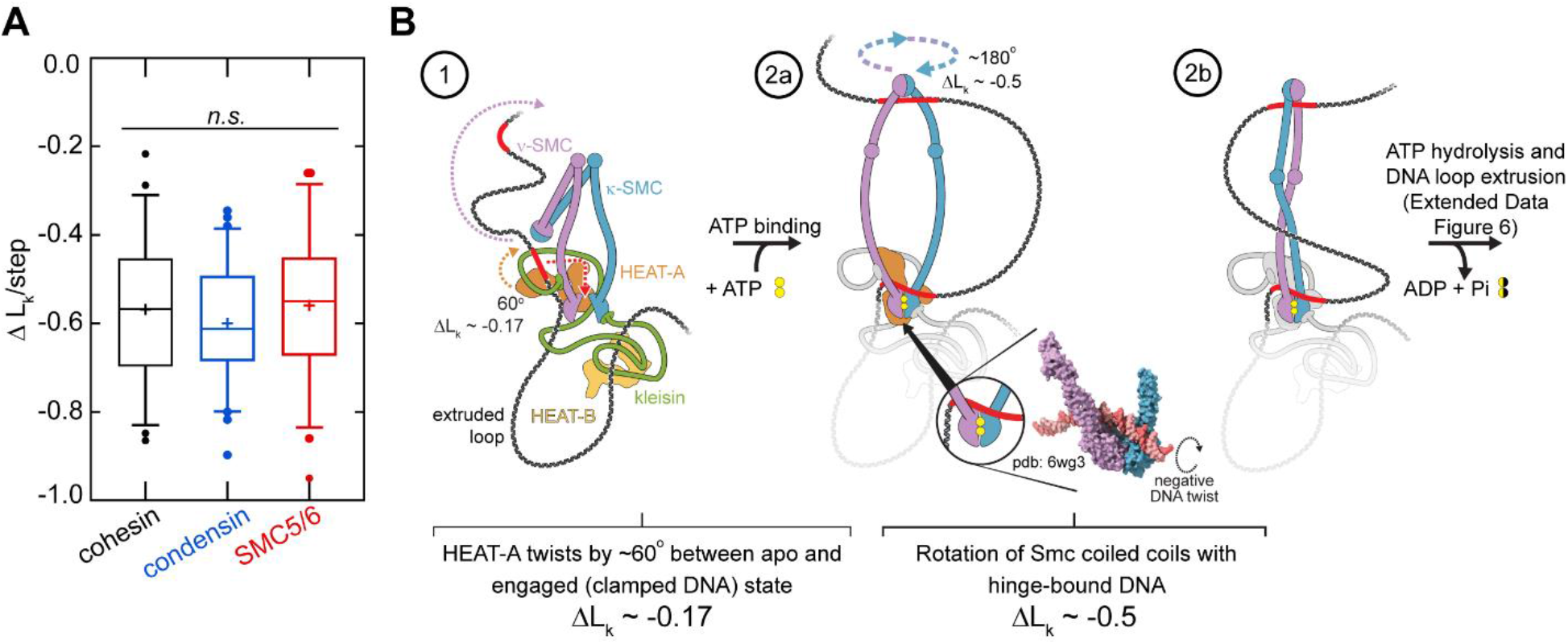
Eukaryotic SMC complexes induce comparable negative DNA twist per loop extrusion step. (**A**) Induced twist per step for all three eukaryotic SMC complexes, where all values of the changes in the linking number Δ*L*_k_ per DNA-loop-extrusion step were pooled from Figs. 2B, 2E, 2H (N = 59 for cohesin, N = 64 for yeast condensin, N = 56 for SMC5/6). Statistical significance was assessed using a one-way analysis of variance (ANOVA) with a significance level *α* = 0.05 (95% confidence interval; n.s. = p > 0.05). (**B**) Potential mechanism yielding a negative DNA twist of Δ*L*_k_ ∼ -0.6 based on the Reel & Seal model (ref. (*19*)). DNA is nontopologicaly held by kleisin and HEAT-A (step 1). Upon ATP binding, clamping of DNA on top of the engaged ATPase heads temporarily forces a small loop of DNA pseudo-topologically into the Smc lumen (step 2a). Rotation of HEAT-A (if bound to DNA in the apo as well as the engaged state) induces a negative twist of about 60°, yielding Δ*L*_k_ ∼-0.17 (Figs. S3D, S3E), into the loop (see inset for twist direction(*43*)). Spontaneous rotation of the coiled coils with hinge-bound DNA by about 180° (ref. (*52–54*)) induces a further linking number change of Δ*L*_k_ ∼-0.5 into the loop (step 2b).

## DISCUSSION

We here presented a high-resolution magnetic tweezers assay on single torsionally constrained DNA molecules that directly measures how much twist is induced by an individual SMC in each loop-extrusion step. The data provide direct evidence that all eukaryotic SMC complexes negatively twist DNA by the same amount at every loop-extrusion step. The linking number change of Δ*L*_k_ = -0.6 is generated upon ATP binding and is found to be a conserved value across the eukaryotic SMC family.

Previous deletion and mutation studies with bulk plasmid assays suggested that SMC proteins induce positive supercoiling of genomic DNA *in vivo* in eukaryotes (*39*) and in prokaryotes (*40, 41*). Positive DNA supercoiling by *Xenopus* 13S condensin was also reconstituted *in vitro*(*20*). Recently, however, it was discovered that two modes of supercoiling can be observed in these plasmid assays for condensing (*25*) and cohesin (Davidson *et al*., 2024, bioRxiv), depending on the molar ratio of SMC to plasmid as well as the absolute protein and DNA concentrations (*25*). In these studies it appears that condensin and cohesin twist DNA positively when it is in large excess to DNA, whereas negative twist is induced at low protein-DNA ratios and absolute concentrations ((*25*) and Davidson et al., 2024, bioRxiv). Our results are in agreement with the negative supercoiling induced at a low concentration of condensing (*25*) and cohesin (Davidson *et a*l., 2024, bioRxiv). In these studies, it remained unclear what constitutes the change in twist direction induced by these SMCs on a molecular level.

Kimura *et al*. (*20*) noted that ATP hydrolysis is necessary for the supercoiling reaction to occur. By contrast, for conditions where ATP hydrolysis was inhibited we observed that SMCs generate at most one step with -0.6 turns (Figs.2A, 2D, 2G, S2) – which shows that SMCs induce negative twist into DNA upon ATP binding alone. Possibly, the topoisomerases used in the experiments of ref. (*20*) was unable to catalyze sufficient changes in linking number to be resolved in these bulk assays. In contrast, Martínez-García *et al*. (*25*) found that incubation of the supercoiling reaction with ATP and subsequent incubation with an excess of AMP-PnP increases the amount of topo I-fixed supercoils 2-fold compared to an incubation with ATP alone. From the average levels of induced supercoiling, the authors estimated that a negative twist of Δ*L*_k_ = -0.8 is induced upon ATP binding which is reduced to Δ*L*_k_ = -0.4 upon ATP hydrolysis. While the induction of negative supercoils during each extrusion step aligns with our observation, the Δ*L*_k_ values differ from our direct single-molecule measurements, as we observed that ATP binding alone induces a twist of Δ*L*_k_ = -0.6 which remains unchanged by subsequent ATP hydrolysis. Possibly, condensin is stabilized on DNA in the ATP-bound state in the plasmid assay, preventing the loss of condensin-mediated supercoils upon loop dissociation before topo I could have permanently induced a linking number change. The discrepancy between the estimates of Δ*L*_k_ per ATPase cycle by Martínez-García *et al*. (*25*) and our results may be explained by the assumptions that had to be made in the bulk assay to assess the supercoiling activity of condensin. Specifically, all condensin molecules were assumed to be active and topo I was assumed to be 100% efficient in removing the generated supercoils. In contrast, our single-molecule experiments directly measure the induced twist on the single-step level in loop extrusion by a single SMC complex, without any treatment by topoisomerases.

When we converted the SMC5/6 hexamer to a SMC5/6 octamer, we observed that the octamer can induce DNA twist in a single step (inducing a linking number change of Δ*L*_k_ = -0.51 ± 0.17, mean ± SD) despite its inability to hydrolyze ATP (*37, 42*) or extrude DNA loops (Figs. 2G, S2C and refs. (*16, 36*)), suggesting that the SMC5/6 octamer is able to bind ATP. Since the SMC5/6 octamer has been shown to topologically embrace DNA (*16, 38*), it is possible that the ATP binding-induced power stroke by the SMC5/6 octamer serves as an intermediate to the topological loading reaction of SMC complexes onto DNA. Current models of DNA loop extrusion postulate that a transient DNA loop is inserted pseudo-topologically into the SMC lumen upon ATP binding to the complex due to a DNA clamping onto the engaged Smc ATPase heads (*19, 38, 43–50*) which causes the power stroke (*45*). Transient hinge opening in this state would lead to a topological entrapment of the DNA duplex (see Fig. S3F-3H for such a potential pathway to topological entrapment).

Since DNA twisting occurs concomitantly with loop extrusion, we conclude that DNA twisting by -0.6 turns in each step is inherent to the ATP-driven DNA loop extrusion cycle of SMC complexes. While the DNA loop extrusion mechanism still remains incompletely resolved, future models have to quantitatively consider the twist of -0.6 turns induced into the loop upon ATP binding. Based on previous structural data, we can point to a few elements that may explain the mechanism driving the induced negative supercoiling (Fig. 3B) in a Reel & Seal model (*19*) with a twist. Upon ATP binding, the HEAT-A subunit (accessory subunit I, NIPBL-Mau2 for human cohesin, in Fig. S1A) undergoes a rigid body rotation about 60° (ΔL_k_ ∼ -0.17) which forms a positively charged channel to clamp DNA onto the ATPase heads (*43, 51*) (Figs. 3B step 1 to 2a, S3A, S3B). While the DNA trajectory within SMC complexes in the apo state has so far not been resolved, continuous DNA binding to HEAT-A in both states may direct DNA to the binding site formed by the engaged ATPase heads and thus induce ΔL_k_ ∼ -0.17 into the extruded loop. Upon head engagement, the coiled coils unfold (*52*) and may spontaneously intertwine as has been observed by negative staining Electron Microscopy (*53*) (Fig. S3C), high-speed Atomic Force Microscopy (*52*) (Fig. S3D), and in Molecular Dynamics simulations (*54*) (Fig. 3B step 2a-b). DNA binding to the hinge has been proposed to be essential for DNA loop extrusion(*52*). If intertwining of the coiled coils occurs after DNA is bound at both the hinge and the ATPase heads, this would induce a negative linking number change of about -0.5 into the DNA loop. Such a mechanism is supported by the finding that a mutant which is defective in DNA binding at the hinge (*52*) shows reduced supercoiling activity (Davidson *et al*., 2024, bioRxiv). These two contributions, the rigid-body rotation of HEAT-A and the spontaneous intertwining of the Smc coiled coils, therefore may additively contribute an induced linking number of -0.6 to -0.7, which is close to the observed negative linking number change of -0.6 upon ATP binding. Subsequent ATP hydrolysis disengages the ATPase heads and transfers the accumulated twist into the extruded loop, clearing the way the next cycle of DNA loop extrusion (Fig. S3E).

What is the impact of SMC-induced DNA supercoiling on the genome? DNA binding of CCCTC-binding factor (CTCF) demarcates topologically associated domains (TADs) *via* an orientated interaction of CTCF with cohesion (*28, 55–61*). The 2- to 3-fold higher contact frequency within TADs than across TADs (*62–65*) might be partially explained by the barrier function of CTCF on loop progression. However, it is also plausible that supercoiling accumulates within a subset of (sub-)TADs (*66*) which could account for the increased intra-TAD contact frequency (*65*). While the accumulation of supercoils within TADs has so far been thought to be due to transcription-mediated supercoiling (*65, 67, 68*), this could also be the result of cohesin-mediated loop extrusion itself. The accumulation of supercoiling within TADs can contribute to their insulating property (*69*) as well as to the facilitation of long-range contacts between genomic loci within a TAD, e.g. promoter-enhancer contacts (*70*). The latter could explain how cohesin may drive transcription of enhancer-controlled developmental genes even in the absence of loop anchors positioned at enhancer-promoter pairs(*10*). SMCs preferentially bind and loop onto positively over negatively supercoiled DNA (*23, 24, 71*), which could prevent SMCs to load onto already extruded (negatively supercoiled) extruded loops. This bias could contribute to the appearance of characteristic CTCF-dependent TAD corner dots in Hi-C maps (*57, 72, 73*) since those depend on a fraction of TADs whose DNA is completely contained in SMC-mediated loops (*74*) as observed *in vivo* (*75, 76*). Similarly, extrusion of entire chromosomes without gaps between loop anchors is instrumental to compact metazoan mitotic chromosomes for sister chromatid segregation (*74*), which could also be mediated by the preferential recruitment to positively supercoiled, non-looped DNA. Similarly, the SMC-mediated positive supercoiling of genomic regions outside loops might direct topo II to decatenate sister chromatids before relaxing supercoils (*39*). If bacterial SMC complexes induce a similar amount of twist as their eukaryotic relatives, this could contribute significantly to the observed supercoiling density of many bacteria *in vivo* (supercoiling density σ ∼-0.06, ref. (*77, 78*)), since we observed that SMCs induce supercoiling densities of -0.04 to -0.06 in the extruded DNA (for example, σ ∼-0.6/150 bp*10.5/bp ∼ -0.04, given a typical step size of ∼150 bp (ref. (*27*) and Fig. S1D)).

Our observation that all eukaryotic SMC complexes twist DNA negatively at every loop-extrusion step establishes DNA supercoiling as an integral part of the DNA loop-extrusion mechanism. The measured quantitative loop extrusion-induced DNA twist of -0.59 ± 0.02 (mean ± SD) will help modelling and polymer simulations to capture yet finer structures of genomes(*65*), and will inform refined models of DNA loop extrusion with the handedness and amount of induced twist upon ATP binding.

## ACKNOWLEDGMENTS

We thank Jaco van der Torre for fruitful discussions. Research in the laboratory of C.D. was funded by ERC Advanced Grant 883684 (DNA looping), NWO grant OCENW.GROOT.2019.012, and the BaSyC program. Research in the laboratory of J.-M.P. was supported by Boehringer Ingelheim, the Austrian Research Promotion Agency (Headquarter grant FFG-FO999902549), the European Research Council under the European Union’s Horizon 2020 research and innovation program GA No 101020558, the Human Frontier Science Program (grant RGP0057/2018) and the Vienna Science and Technology Fund (grant LS19-029). Research in the laboratory of S.G. was supported by ERC Consolidator Grant 724482.

## AUTHOR CONTRIBUTIONS

R.J., R.B., and C.D. designed experiments; R.J and R.B. conducted single-molecule experiments; R.J. analysed the single-molecule data; R.B. made structure predictions; I.D. purified human cohesin; M.T. purified yeast SMC5/6; R.B., R.J, and C.D. wrote the manuscript; S.G., J.-M.P., and C.D. acquired funding and supervised the work.

## COMPETING INTERESTS

Authors declare that they have no competing interests.

## DATA AND MATERIAL AVAILABILITY

All data are available in the main text or the supplementary materials.

## SUPPLEMENTARY MATERIALS

Materials and Methods

Figs. S1 to S3

## Notes

### Competing Interest Statement

The authors have declared no competing interest.

## REFERENCES AND NOTES

1. S. H. Harvey, M. J. Krien, M. J. O’Connell, Genome Biol., in press, doi:10.1186/gb-2002-3-2-reviews3003.

2. E. Kim, R. Barth, C. Dekker, Annu. Rev. Biochem. 92, 15–41 (2023).

3. C. H. Haering, A. M. Farcas, P. Arumugam, J. Metson, K. Nasmyth, Nature. 454, 297–301 (2008).

4. J.-M. Peters, T. Nishiyama, Cold Spring Harb. Perspect. Biol. 4, a011130–a011130 (2012).

5. M. Ganji et al., Science. 360, 102–105 (2018).

6. J. H. Gibcus et al., Science. 359 (2018), doi:10.1126/science.aao6135.

7. E. P. Nora et al., Nat. Commun. 11, 5612 (2020).

8. I. F. Davidson et al., Science. 366, 1338–1345 (2019).

9. I. F. Davidson, J. M. Peters, Nat. Rev. Mol. Cell Biol. 22, 445–464 (2021).

10. J. A. Horsfield, FEBS J., in press, doi:10.1111/febs.16362.

11. A. Piazza et al., Nat. Cell Biol. 23, 1176–1186 (2021).

12. I. Litwin, E. Pilarczyk, R. Wysocki, Genes. 9, 581 (2018).

13. P. Batty et al., EMBO J. 42, e113475 (2023).

14. M. I. Fousteri, EMBO J. 19, 1691–1702 (2000).

15. Y. Kim, Z. Shi, H. Zhang, I. J. Finkelstein, H. Yu, Science. 366, 1345–1349 (2019).

16. B. Pradhan et al., Nature. 616, 843–848 (2023).

17. S. Golfier, T. Quail, H. Kimura, J. Brugués, eLife. 9, 1–34 (2020).

18. M. Kong et al., Mol. Cell. 79, 99–114.e9 (2020).

19. C. Dekker, C. H. Haering, J.-M. Peters, B. D. Rowland, Science. 382, 646–648 (2023).

20. K. Kimura, T. Hirano, Cell. 90, 625–634 (1997).

21. K. Kimura, V. V. Rybenkov, N. J. Crisona, T. Hirano, N. R. Cozzarelli, Cell. 98, 239–248 (1999).

22. K. A. Hagstrom, V. F. Holmes, N. R. Cozzarelli, B. J. Meyer, Genes Dev. 16, 729–742 (2002).

23. E. Kim, A. M. Gonzalez, B. Pradhan, J. van der Torre, C. Dekker, Nat. Struct. Mol. Biol. (2022), doi:10.1038/s41594-022-00802-x.

24. K. Jeppsson et al., Mol. Cell, S1097276524000066 (2024).

25. B. Martínez-García et al., EMBO J. 42, e111913 (2023).

26. K. Krassovsky, R. P. Ghosh, B. J. Meyer, Genome Res. 31, 1187–1202 (2021).

27. J.-K. Ryu et al., Nucleic Acids Res. 50, 820–832 (2022).

28. I. F. Davidson et al., Nature. 616, 822–827 (2023).

29. H. Dohnalová et al., “Temperature-dependent twist of double-stranded RNA probed by magnetic tweezers experiments and molecular dynamics simulations” (preprint, Biophysics, 2023),, doi:10.1101/2023.05.31.543084.

30. T. R. Strick, J.-F. Allemand, D. Bensimon, A. Bensimon, V. Croquette, Science. 271, 1835–1837 (1996).

31. F. Mosconi, J. F. Allemand, D. Bensimon, V. Croquette, Phys. Rev. Lett. 102, 078301 (2009).

32. R. Vlijm, A. Mashaghi, S. Bernard, M. Modesti, C. Dekker, Nanoscale. 7, 3205–3216 (2015).

33. E. Ostrofet, F. S. Papini, D. Dulin, Sci. Rep. 8, 15920 (2018).

34. S. Forth et al., Phys. Rev. Lett. 100, 148301 (2008).

35. T. Lionnet et al., Nucleic Acids Res. 34, 4232–4244 (2006).

36. R. Barth et al., “SMC motor proteins extrude DNA asymmetrically and contain a direction switch” (preprint, Molecular Biology, 2023),, doi:10.1101/2023.12.21.572892.

37. M. Taschner et al., EMBO J. 40, e107807 (2021).

38. M. Taschner, S. Gruber, Nat. Struct. Mol. Biol. 30, 619–628 (2023).

39. J. Baxter et al., Science. 331, 1328–1332 (2011).

40. J. C. Lindow, R. A. Britton, A. D. Grossman, J. Bacteriol. 184, 5317–5322 (2002).

41. J. A. Sawitzke, S. Austin, Proc. Natl. Acad. Sci. 97, 1671–1676 (2000).

42. S. T. Hallett et al., Nucleic Acids Res. 49, 4534–4549 (2021).

43. Z. Shi, H. Gao, X. C. Bai, H. Yu, Science. 368, 1454–1459 (2020).

44. I. A. Shaltiel et al., Science. 376, 1087–1094 (2022).

45. S. K. Nomidis, E. Carlon, S. Gruber, J. F. Marko, Nucleic Acids Res. 50, 4974–4987 (2022).

46. J. F. Marko, P. De Los Rios, A. Barducci, S. Gruber, Nucleic Acids Res. 47, 6956–6972 (2019).

47. M. L. Diebold-Durand et al., Mol. Cell. 67, 334–347.e5 (2017).

48. Y. Yu et al., 1–10 (2022).

49. B. G. Lee, J. Rhodes, J. Löwe, Proc. Natl. Acad. Sci. U. S. A. 119, 1–10 (2022).

50. J. E. Collier et al., eLife. 9, 1–36 (2020).

51. T. L. Higashi et al., Mol. Cell. 79, 917–933.e9 (2020).

52. B. W. Bauer et al., Cell. 184, 5448–5464 (2021).

53. M. T. Hons et al., Nat. Commun. 7, 12523 (2016).

54. D. Krepel, A. Davtyan, N. P. Schafer, P. G. Wolynes, J. N. Onuchic, Proc. Natl. Acad. Sci. 117, 1468–1477 (2020).

55. E. P. Nora et al., Cell. 169, 930–944.e22 (2017).

56. G. Wutz et al., EMBO J. 36, 3573–3599 (2017).

57. S. S. P. Rao et al., Cell. 159, 1665–1680 (2014).

58. J. R. Dixon et al., Nature. 485, 376–380 (2012).

59. Y. Li et al., Nature. 578, 472–476 (2020).

60. G. Fudenberg et al., Cell Rep. 15, 2038–2049 (2016).

61. A. L. Sanborn et al., Proc. Natl. Acad. Sci. U. S. A. 112, E6456–E6465 (2015).

62. E. H. Finn et al., Cell. 176, 1502–1515.e10 (2019).

63. A. S. Hansen, C. Cattoglio, X. Darzacq, R. Tjian, Nucleus. 9, 20–32 (2018).

64. L. H. Chang, S. Ghosh, D. Noordermeer, J. Mol. Biol. 432, 643–652 (2020).

65. D. Racko, F. Benedetti, J. Dorier, A. Stasiak, Nucleic Acids Res. 47, 521–532 (2019).

66. C. Naughton et al., Nat. Struct. Mol. Biol. 20, 387–395 (2013).

67. D. Racko, F. Benedetti, J. Dorier, A. Stasiak, Nucleic Acids Res. 46, 1648–1660 (2018).

68. R. Janissen, R. Barth, M. Polinder, J. Van Der Torre, C. Dekker, “Single-molecule visualization of twin-supercoiled domains generated during transcription” (preprint, Biophysics, 2023),, doi:10.1101/2023.08.25.554779.

69. F. Benedetti, J. Dorier, Y. Burnier, A. Stasiak, Nucleic Acids Res. 42, 2848–2855 (2014).

70. F. Benedetti, J. Dorier, A. Stasiak, Nucleic Acids Res. 42, 10425–10432 (2014).

71. A. Diman et al., “Human Smc5/6 recognises transcription-generated positive DNA supercoils” (preprint, Molecular Biology, 2023),, doi:10.1101/2023.05.04.539344.

72. Z. Tang et al., Cell. 163, 1611–1627 (2015).

73. E. de Wit et al., Mol. Cell. 60, 676–684 (2015).

74. . J. Banigan, A. A. van den Berg, H. B. Brandão, J. F. Marko, L. A. Mirny, eLife. 9, e53558 (2020).

75. M. Gabriele et al., Science. 376, 476–501 (2022).

76. P. Mach et al., Nat. Genet. 54, 1907–1918 (2022).

77. W. R. Bauer, Annu. Rev. Biophys. Bioeng. 7, 287–313 (1978).

78. E. L. Zechiedrich et al., J. Biol. Chem. 275, 8103–8113 (2000).

79. J. Lipfert, J. W. J. Kerssemakers, T. Jager, N. H. Dekker, Nat. Methods. 7, 977–980 (2010).

80. R. Janissen et al., Nucleic Acids Res., gku677. (2014).

81. R. Janissen, B. Eslami-Mossallam, I. Artsimovitch, M. Depken, N. H. Dekker, Cell Rep. 39, 110749 (2022).

82. J. P. Cnossen, D. Dulin, N. H. Dekker, Rev. Sci. Instrum. 85, 103712 (2014).

83. T. D. Goddard et al., Protein Sci. 27, 14–25 (2018).

84. N. J. Petela et al., eLife. 10, 1–32 (2021).

85. M. Srinivasan et al., Cell. 173, 1508–1519.e18 (2018).

86. D. Klaue, R. Seidel, Phys. Rev. Lett. 102 (2009), doi:10.1103/PhysRevLett.102.028302.

87. L. Loeff, J. W. J. Kerssemakers, C. Joo, C. Dekker, Patterns. 2, 100256 (2021).

88. J.-K. Ryu et al., Nucleic Acids Res. 50, 820–832 (2022).

89. J. E. Collier, K. A. Nasmyth, eLife. 11, e80310 (2022).

90. S. Gruber et al., Cell. 127, 523–537 (2006).

91. J. Buheitel, O. Stemmann, EMBO J. 32, 666–676 (2013).

92. K. Nasmyth, Nat. Cell Biol. 13, 1170–1177 (2011).

93. K. Nagasaka et al., Mol. Cell. 83, 3049–3063.e6 (2023).

94. L. Wilhelm et al., eLife. 4 (2015), doi:10.7554/eLife.06659.

95. G. Ö. Çamdere, K. K. Carlborg, D. Koshland, Proc. Natl. Acad. Sci. 115, 9732–9737 (2018).

96. M. Minamino, T. L. Higashi, C. Bouchoux, F. Uhlmann, Life Sci. Alliance. 1, e201800143 (2018).

97. Y. Murayama, F. Uhlmann, Nature. 505, 367–371 (2014).

98. M. Hassler et al., Mol. Cell. 74, 1175–1188.e9 (2019).

