## Supplementary material for "All eukaryotic SMC proteins induce a twist of -0.6 at each DNA-loop-extrusion step": Methods and Supplemental Figures

Science AAAS

**The PDF file includes:**

Materials and Methods
Figs. S1 to S3

### Materials and Methods

Protein expression and purification

WT and EQ/EQ human cohesin, as well as NIPBL-Mau2 were expressed in and purified from *Sf9* insect cells as described previously (*8*). *S. cerevisiae* condensin was expressed in and purified from *S. cerevisiae* as described previously (*5*). *S. cerevisiae* SMC5/6 was expressed in and purified from *E. coli* as described previously (*38*).

Synthesis of torsionally constrained dsDNA

Linear, torsionally constrained dsDNA constructs were synthesized based on a 3.6 kbp fragment of the pRL-SV40 plasmid (Promega, USA), digested with BamHI and XbaI. Afterwards, the fragment was enzymatically ligated to 600 bp dsDNA handles that contained multiple digoxigenin on one end of the dsDNA construct and multiple biotin at the other end of the dsDNA construct, as previously described (*79, 80*). To synthesize these handles, a 1.2 kb fragment from pBluescript (Stratagene, USA) was amplified by PCR in the presence of Biotin-16-dUTP (Roche, Switzerland) and Digoxigenin-11-dUTP (Roche, Switzerland) in a 1:10 ratio, using forward primer 5'-GACCGAGATAGGGTTGAGTG and reverse primer 5'-CAGGGTCGGAACAGGAGAGC. Prior to enzymatic ligation *via* T4 DNA ligase (New England Biolabs, UK) overnight, the handles fragment was digested with either BamHI or XbaI. The final dsDNA construct was cleaned up from the access of handles by running on a 1% agarose gel and extract the dsDNA construct using a gel purification kit (A9282, Promega).

Magnetic tweezers

The magnetic tweezers used in this study was described previously (*27, 81*). Briefly, light transmitted through the sample was collected by a 50x oil-immersion objective (CFI Plan 50XH, Achromat; 50x; NA = 0.9, Nikon) and projected onto a 4-megapixel CMOS camera (4M60, Falcon2; Teledyne Dalsa) with a sampling frequency of 60 Hz. The applied magnetic field was generated by a pair of vertically aligned permanent neodymium-iron-boron magnets (Supermagnete GmbH, Germany), separated by a distance of 1 mm, and suspended on a motorized stage (M-126.PD2, Physik Instrumente) above the flow cell. In addition, the magnet pair could be rotated around the illumination axis by an applied DC servo step motor (C-150.PD; Physik Instrumente). Image processing of the collected light allowed to track the real-time position of both surface attached reference beads and superparamagnetic beads coupled to the torsionally constrained dsDNA construct in three dimensions over time. Bead x, y, z position tracking was achieved with a spatial resolution of ~2 nm (*27*)) using a cross-correlation algorithm realized with custom-written software in LabView (2011, National Instruments Corporation) (*82*). The software determined the bead positions with spectral corrections to correct for camera blur and aliasing.

Measurement of DNA loop extrusion-induced DNA twist

The flow cell preparation for the magnetic tweezers experiments used in this study has been described previously in detail (*27, 81*). Briefly, polystyrene reference beads (Polysciences Europe) of 1.5 µm in diameter were diluted 1:1500 in PBS buffer (pH 7.4) and adhered to 1 M KOH treated surface of the flow cell channel as fiducial markers. Next, 0.5 mg/ml digoxigenin sheep antibody Fab fragments (Roche, Switzerland) in PBS buffer were incubated for 1 h within the flow cell channel, following incubation for 2 h of 10 mg/ml BSA (New England Biolabs, UK), diluted in PBS buffer. For each experiment, 1 pM of the torsionally constrained dsDNA construct was incubated in PBS buffer for 20 min in the flow cell channel. After washing with 500 ml PBS buffer, the addition of 100 µl streptavidin-coated superparamagnetic beads with a diameter of 1 µm (diluted 1:400 in PBS buffer; MyOne #65601 Dynabeads, Invitrogen/Life Technologies) for 5 min resulted in the attachment of the beads to biotinylated dsDNA construct. Non-attached beads were washed out with PBS buffer. Prior to conducting the force-extension experiments, we assessed whether dsDNA tethers were singly tethered by applying a high force (5 pN) and 20 negative rotations, and if they were torsionally constrained by applying 20 rotations to each direction at low force (0.3 pN). Only single and torsionally constrained DNA tethers were analyzed. All experiments were conducted at 22.3°C using a buffer containing 40 mM Tris pH 7.5, 40 mM NaCl, 2.5 mM $MgCl_2$, 1 mM DTT, 1 mM ATP, 0.05% Tween-20, 0.25 mg/ml BSA.

For assessing the twist generated during the DNA loop extrusion experiments, a reference rotation curve was first acquired at 0.3 pN with negative and positive rotations of each 12 turns, with a rotation speed of 0.25 turns $s^{-1}$. Subsequently, the SMC proteins (Cohesin WT: 20-100 pM; Cohesin EQ/EQ: 10-20 pM; Condensin: 0.4-1 nM; SMC5/6: 3-6 nM) in presence of 1 mM ATP or 1 mM AMP-PnP (Sigma, Netherlands) - as indicated in main text - were inserted into the flow cell channel while applying 7 pN to the dsDNA tethers. The force was then quickly lowered to a constant force of 0.3 pN to record the DNA loop extrusion steps *via* the change in bead Z-position(*27*, *28*) for 10 minutes , which is a time where the DNA loop extrusion-activity of most SMCs ceased. To assess the degree of DNA twist that was introduced by each SMC during DNA loop extrusion, another rotation curve was recorded at 0.3 pN with negative and positive rotations of 12 turns with a rotation speed of 0.25 turns $s^{-1}$.

Modelling of experimentally determined cryo-EM map of apo *S. cerevisiae* cohesin

A full pseudo-atomic model of *S. cerevisiae* cohesin in the apo state (in the absence of ATP and DNA) was fitted into the cryo-EM map EMD-12880 using UCSF ChimeraX (*83*). The lower part containing the ATPase heads of Smc1 and Smc3, kleisin, as well as Scc2 was taken from pdb entry 6ZZ6 (*50*) and the folded coiled coils were taken from pdb entry 7OGT (*84*) as done previously (*85*). The EM maps and protein structures in Figure 3b and Figure S3 were depicted using UCSF ChimeraX (*83*).

Quantification and Statistical Analyses

Magnetic tweezers datasets were processed and analyzed using custom-written Igor v6.37-based scripts. From our raw data, we removed traces that showed surface-adhered magnetic beads as well as dsDNA tethers where the DNA-bead attachment points were close (<100 nm) to the magnetic equator of the magnetic beads using a previously described method (*86*). Tethers that detached from the surface during the measurement were also rejected from further analysis. All traces and rotation curves resulting from experiments conducted at identical conditions for each SMC were pooled and filtered to 1 Hz (moving average) for extrusion-step identification and prior to Gaussian fitting of the rotations curves.

DNA loop extrusion steps sizes were extracted from the change in bead Z-positions in the traces using a previously described methodology and stepfinder algorithm (*27*, *87*). The induced DNA twist during loop extrusion was determined by fitting a Gaussian to the bead Z-positions as a function of applied turns. The peak position of the fitted Gaussian before DNA loop extrusion $C_{\text{before}}$ were subtracted from the peak position determined after the DNA loop extrusion experiments $C_{\text{after}}$ which reflect the induced DNA twists $\Delta L_k = \Delta Tw = C_{\text{after}} - C_{\text{before}}$ in the unit of *turns* for each trace.

The statistical analyses comparing the DNA twist results for each SMC and condition were conducted using one-way analysis of variance (ANOVA) with a significance level $\alpha = 0.05$ (95% CI).

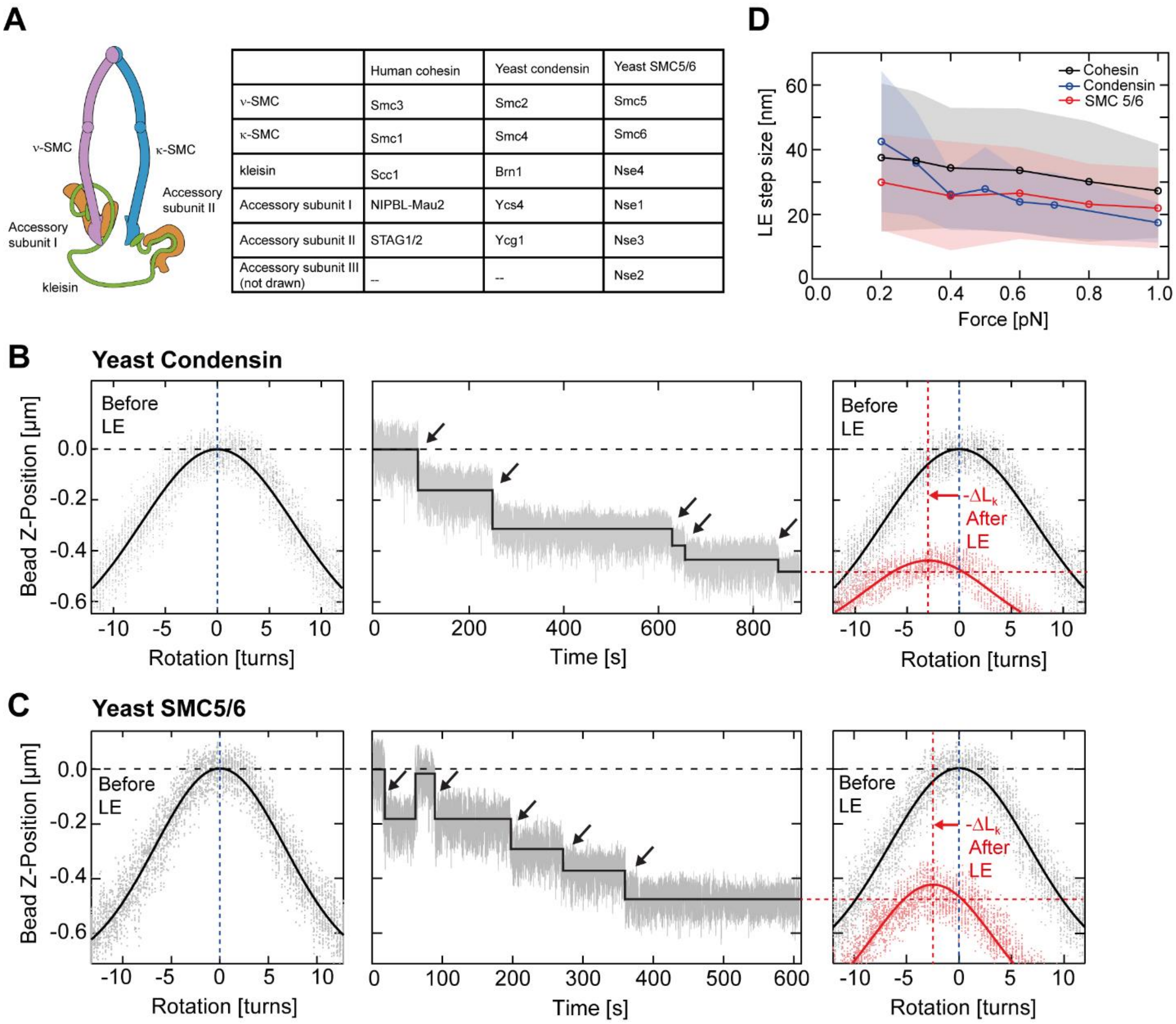


| | Human cohesin | Yeast condensin | Yeast SMC5/6 |
|---|---|---|---|
| ν-SMC | Smc3 | Smc2 | Smc5 |
| κ-SMC | Smc1 | Smc4 | Smc6 |
| kleisin | Scc1 | Brn1 | Nse4 |
| Accessory subunit I | NIPBL-Mau2 | Ycs4 | Nse1 |
| Accessory subunit II | STAG1/2 | Ycg1 | Nse3 |
| Accessory subunit III (not drawn) | -- | -- | Nse2 |

**Fig. S1. Nomenclature and step size distribution versus applied force for human cohesin, yeast condensin, and yeast SMC5/6. Exemplary rotation curves for yeast condensin and yeast SMC5/6. Related to Figs. 1 and 2.**
(**A**) Nomenclature for human cohesin, budding yeast condensin, and budding yeast SMC5/6. (**B**) Same as shown in Figures 1A-1C, for yeast condensin. Black arrows in DNA loop extrusion trajectories depict single steps. (**C**) Same as (B) for yeast SMC5/6. (**D**) DNA loop extrusion step size versus DNA tension for all SMC families. Shown is the median step size as circles. Shaded areas denote the interquartile range as an indication of the spread in the data. The step size data for Condensin originates from the previous study Ryu *et al.* (ref. (*88*)). Sample size (individual steps) per force for human cohesin (black) was N = 959, 1054, 1256, 1430, 1951, 701 for increasing force; for yeast condensin (blue) N = 976, 565, 1344, 468, 605, 510, 556; and for yeast SMC5/6 (red) N = 109, 189, 206, 278, 281.

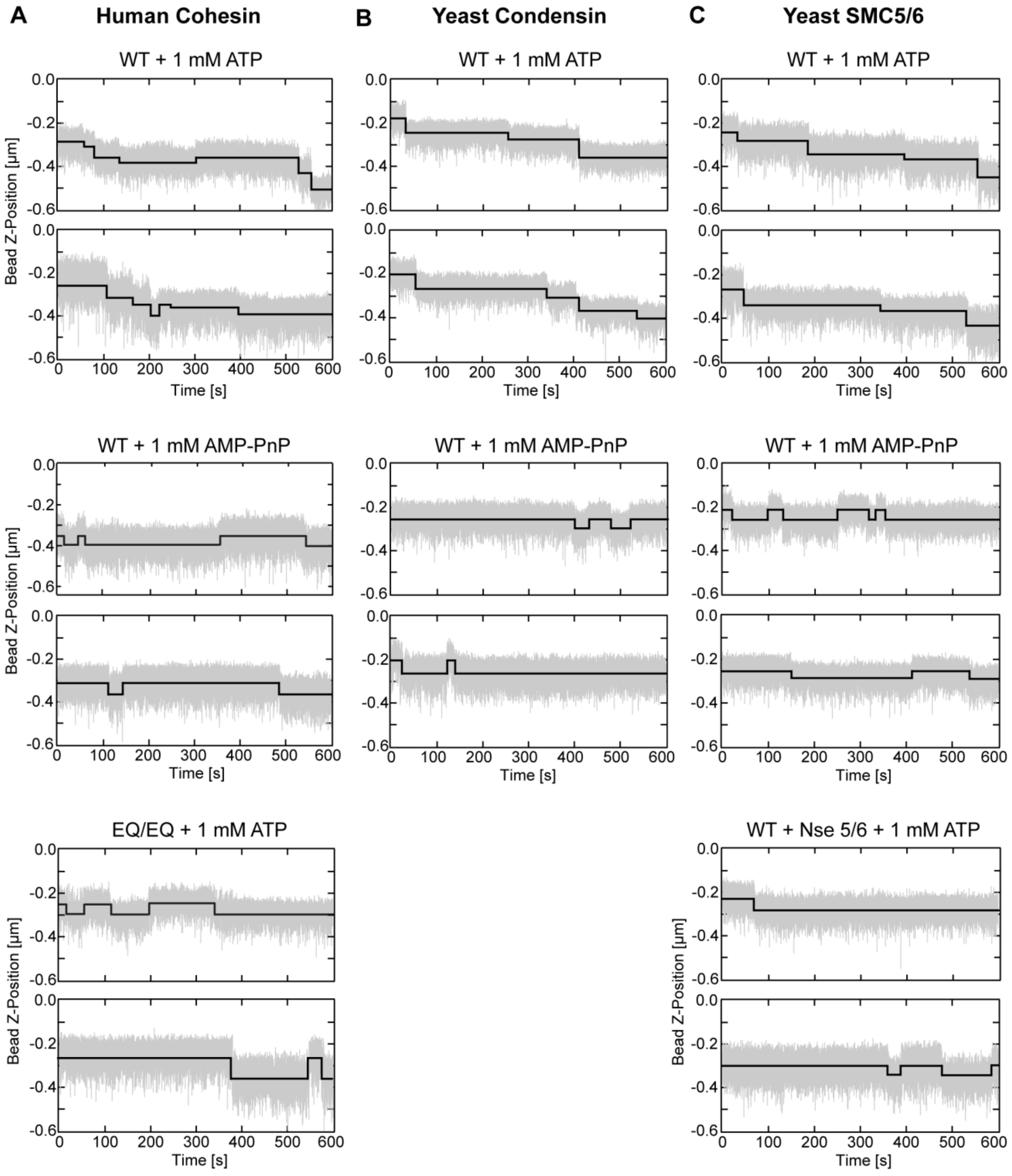


**Fig. S2. Example DNA loop extrusion stepping trajectories for all eukaryotic SMC complexes. Related to Figs. 1 and 2.**

(**A**) Exemplary traces of cohesin-mediated DNA loop extrusion steps in the presence of ATP and WT cohesin, AMP-PnP and WT cohesin, and ATP with the cohesin EQ/EQ mutant. (**B**) Exemplary traces for yeast condensin as in panel (A). (**C**) Exemplary traces for yeast SMC5/6 as in panel (A). Note on the bottom right panel: the SMC5/6 hexamer forms an octamer with Nse5/6. Traces show only a single downward step or repeated downward-reverse steps in the presence of AMP-PNP, the cohesin EQ/EQ mutant with ATP, or the SMC5/6 octamer in the presence of ATP.

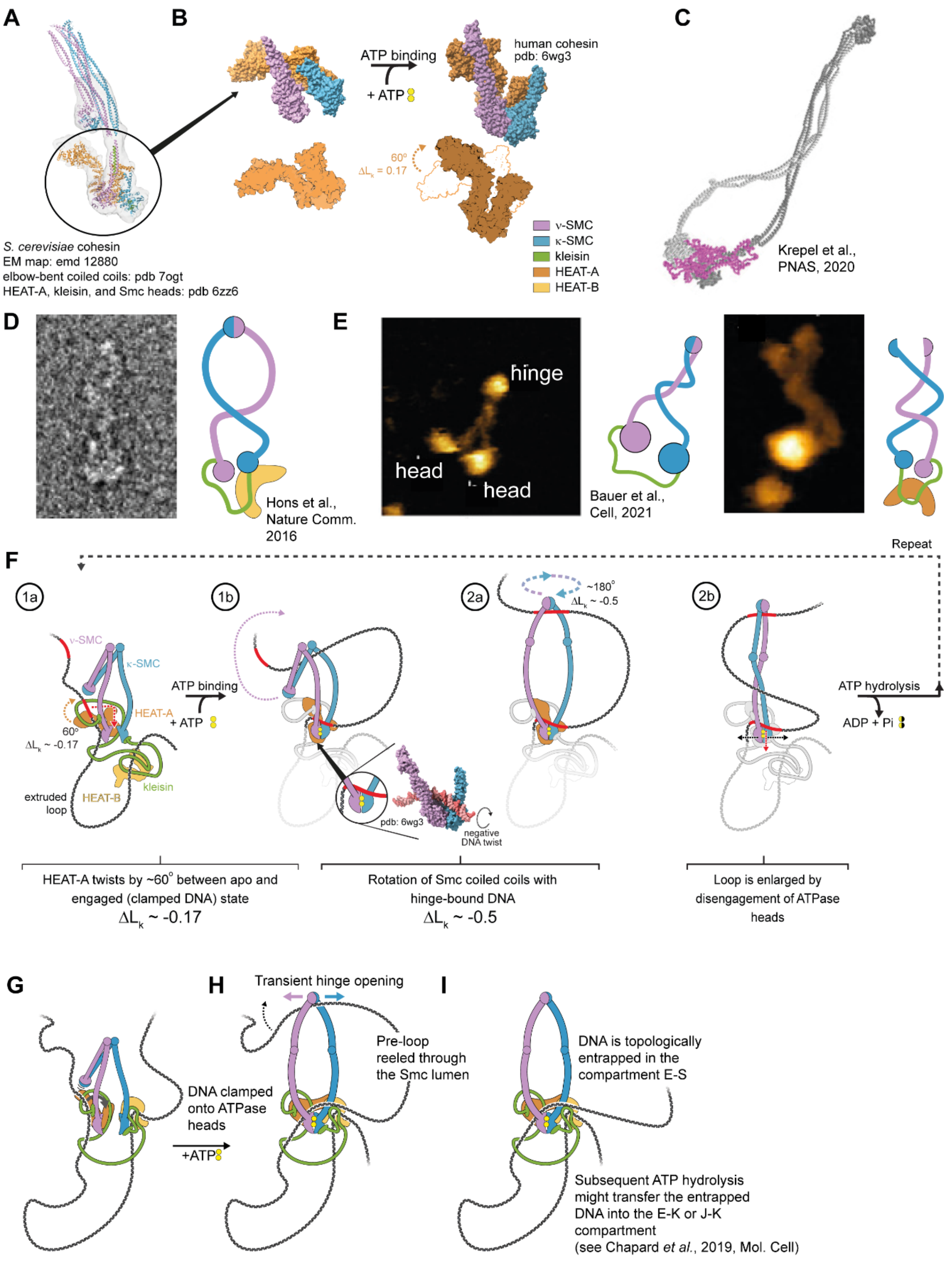
A
B
C
S. cerevisiae cohesin
EM map: emd 12880
elbow-bent coiled coils: pdb 7ogt
HEAT-A, kleisin, and Smc heads: pdb 6zz6
ATP binding
+ ATP
human cohesin
pdb: 6wg3
60°
ΔLk = 0.17
ν-SMC
κ-SMC
kleisin
HEAT-A
HEAT-B
Krepel et al., PNAS, 2020
D
Hons et al., Nature Comm. 2016
E
hinge
head
head
Bauer et al., Cell, 2021
F
Repeat
1a
1b
2a
2b
ν-SMC
κ-SMC
HEAT-A
60°
ΔLk ~ -0.17
extruded loop
HEAT-B
kleisin
ATP binding
+ ATP
pdb: 6wg3
negative DNA twist
~180°
ΔLk ~ -0.5
ATP hydrolysis
ADP + Pi
HEAT-A twists by ~60° between apo and engaged (clamped DNA) state
ΔLk ~ -0.17
Rotation of Smc coiled coils with hinge-bound DNA
ΔLk ~ -0.5
Loop is enlarged by disengagement of ATPase heads
G
H
I
Transient hinge opening
DNA clamped onto ATPase heads
+ATP
Pre-loop reeled through the Smc lumen
DNA is topologically entrapped in the compartment E-S
Subsequent ATP hydrolysis might transfer the entrapped DNA into the E-K or J-K compartment
(see Chapard et al., 2019, Mol. Cell)

**Fig. S3. Structural basis for twists generation upon ATP binding, potential DNA loop extrusion mechanism including twist generation, and potential pathway to topological entrapment upon ATP binding. Related to Fig. 3.**

(**A**) Superimposition of a density map from cryo-EM data for *S. cerevisiae* cohesin(*84*) (EMD-12880) and a pseudo-atomic model in the apo state. (**B**) Rotation of HEAT-A between the pseudo-atomic model in the apo state shown in (a) and the ATP-bound head-engaged gripping state (pdb entry 6wg3; ref. (*43*)). Apo and engaged structures were aligned on the Smc3 ATPase head. HEAT-A rotates by ~ -60$^o$ ($\Delta L_k$ ~ -0.17). (**C**) Conformation of human cohesin as observed by negative staining EM showing intertwined coiled coils, from ref. (*53*). (**D**) Snapshots of human cohesin imaged by high-speed AFM anecdotally showing intertwined coiled coils, adapted from ref. (*52*). (**E**) Reel & Seal model(*19*) with a twist, incorporating an ATP binding-induced negative DNA twist into the loop. DNA is held by kleisin and HEAT-A (step 1a). Clamping of DNA on top of the engaged ATPase heads forces a small loop pseudo-topologically into the Smc lumen (step 1b). The inset to step 1b shows the twisting of the DNA which yields a negatively supercoiled loop (based on pdb entry 6wg3 (ref. (*43*))). Rotation of HEAT-A (if bound to DNA in the apo as well as the engaged state) induces a negative twist (see panels (A-B)). Spontaneous rotation of the coiled coils with hinge-bound DNA by ~180$^o$ (ref. (*52–54*)) induces a further linking number change of $\Delta L_k$ ~ -0.5 into the loop (step 2a-b). ATP hydrolysis causes disengagement of the heads, merging pre-loop and already extruded loop *via* HEAT-A-kleisin latch opening. (**F**) A potential pathway to topological entrapment upon ATP binding. DNA is held at the n-SMC and by accessory subunit I. (**G**) Upon ATP binding and subsequent head engagement, DNA binding on top of the ATPase heads inserts a short loop of DNA (of the length of one SMC-mediated loop extrusion step) within the loop formed by the Smc arms and the engaged Smc heads (E-S compartment). Hinge opening in this state expels the head-bound DNA segment from the Smc lumen, leading to a topological entrapment within the E-S compartment. (**H**) A single duplex of DNA is topologically embraced in the E-S compartment. Subsequent ATP hydrolysis causes the heads to disengage. The DNA is then in the S-K compartment and may be transferred to the E-K compartment in the subsequent ATP binding and head engagement. The sketched pathway is consistent with the fact that the hinge is the main DNA entry gate for topological entrapment(*85, 89–92*) and that a cohesin hinge mutant can perform loop extrusion but is defective in cohesion establishment(*93*). Furthermore, it is in line with the fact that ATP binding, and thus the head engagement, is sufficient to allow topological entrapment(*50, 94–96*) and that clamping of the DNA onto the engaged ATPase heads by HEAT-A is required for topological entrapment in order to insert a transient loop into the E-S compartment. (**I**) Subsequent ATP hydrolysis may transfer the entrapped DNA into the S-K compartment. Note that for SMC5/6 the neck gate (the n-SMC-kleisin interface) appears to be the major or only DNA entry gate(*38*), while DNA entry though the neck gate as well as the hinge can be observed for cohesin and condensin(*51, 85, 89–92, 97, 98*). Topological entry of DNA through the hinge is thus not the only possible entrapment mechanism upon ATP binding since an ATP binding-induced power stroke has also been proposed to enable topological DNA loading thorough the neck gate(*38*).
